# Rewiring glucose metabolism improves 5-FU efficacy in glycolytic p53-deficient colorectal tumors

**DOI:** 10.1101/2021.11.11.468185

**Authors:** Marlies C. Ludikhuize, Sira Gevers, Nguyen T.B. Nguyen, Maaike Meerlo, S. Khadijeh Shafiei Roudbari, M. Can Gulersonmez, Edwin C.A. Stigter, Jarno Drost, Hans Clevers, Boudewijn M.T. Burgering, Maria J. Rodríguez Colman

## Abstract

5-fluorouracil (5-FU) is the backbone for chemotherapy in colorectal cancer (CRC). Response rates in patients are, however, limited to 50%. Despite the importance of 5-FU, the molecular mechanisms by which it induces toxicity remain unclear, limiting the development of strategies to improve efficacy. How fundamental aspects of cancer, such as driver mutations and phenotypic intra-tumor heterogeneity, relate to the 5-FU response is also ill-defined. This is largely due to the shortage of mechanistic studies executed in pre-clinical models that can faithfully recapitulate key CRC features. Here, we analyzed the 5-FU response in human organoids genetically engineered to reproduce the different stages of CRC progression. We find that 5-FU induces pyrimidine imbalance, which leads to DNA damage and cell death. Actively proliferating cancer (stem) cells are accordingly efficiently targeted by 5-FU. Importantly, p53 behaves as a discriminating factor for 5-FU sensitivity, whereas p53-deficiency leads to DNA damage-induced cell death, active p53 protects from these effects through inducing cell cycle arrest. Moreover, we find that targeting the Warburg effect, by rewiring glucose metabolism, enhances 5-FU toxicity by further altering the nucleotide pool and without increasing toxicity in healthy-non-transformed cells. Thus, targeting cancer metabolism in combination with replication stress-inducing chemotherapies emerges as a promising strategy for CRC treatment.

## Introduction

Worldwide, colorectal cancer (CRC) is the third most commonly diagnosed type of cancer and the second leading cause of cancer-related mortality^1^. Surgery remains currently the only curative treatment in patients with early stage CRC or with resectable metastases. CRC patients additionally receive adjuvant chemotherapy and patients with unresectable, metastatic CRC entirely rely on chemotherapy^2^. Although 5-FU based chemotherapies have a poor tumor response (rates up to 50%)^4–6^ and do not effectively extent the disease-free survival^2,4,5,7,8^, it remains the most common treatment for CRC (reviewed in ^3^). Furthermore, it remains unclear for which patients 5-FU therapy is beneficial.

Despite the importance of 5-FU, the underlying mechanism of its toxicity is still debated. Upon cellular uptake, 5-FU is converted into active fluorinated metabolites. 5-FUTP and 5-FdUTP can be incorporated into RNA and DNA respectively, and F-dUMP can inhibit thymidylate synthase (TS), impairing the deoxynucleotide pool and consequently DNA replication and repair (reviewed in^3,9^). How this recapitulates in patients remains unclear, although a number of studies show that 5-FU can induce cytotoxicity via F-UTP incorporation into RNA^10–14^. TS expression in the tumor appears, on the other hand, to correlate with the 5-FU response, suggesting that 5-FU’s toxicity could rely on impaired DNA replication and/or repair, however correlation does not imply causation^15–18^, reviewed in^3^.

Although precision medicine and application of targeted therapies have been facilitated by genomic studies, for conventional chemotherapies this has been relatively unsuccessful^2,19^. The relevance of different genetic mutations in determining 5-FU response, in fact, still remains elusive. Furthermore, genetic and phenotypic intra-tumor heterogeneity also contributes to differential therapy response and resistance^2,20–22^. Cancer stem cells (CSCs) are such a subset of tumor cells within the tumor that actively proliferate and exhibit differentiation potential^23–25^. Whether different CRC cell types respond differently to 5-FU also remains to be elucidated. Altogether, there is still a lack of knowledge that limits improvement of 5-FU-based CRC treatment strategies.

Studies aimed to increase our knowledge on the mechanisms of action of conventional chemotherapies should be performed in pre-clinical models that allow manipulation, but at the same time recapitulate the morphological and molecular characteristics of CRC tumors. Tumor-derived 2D cell lines have greatly contributed to the current understanding of cancer biology, but in most cases exhibit poor (genetic) stability and lack the heterogeneity characteristic of tumors *in vivo*^26^. In contrast, tumor-biopsies that exhibit all features of the tumor, fall oftentimes short for chemotherapy response studies due to the limited material and these lack options for manipulation. Patient-derived organoids bridge the gap between patient-biopsies and 2D cell lines. Patient-derived organoids recapitulate somatic copy number variations and mutation spectra found in CRC tumors and the genetic and non-genetic heterogeneity of CRC tumors^27,28^. Moreover, multiple studies have shown that tumor-derived organoids from multiple cancer types predict the chemotherapy response in patients^29–33^ and the response is stable over time^32,34^. Furthermore, organoids allow manipulation to assess causality and the mechanisms downstream drug toxicity.

A hurdle in identifying the role of specific mutations in chemotherapy response is the fact that tumors have a complex genetic background and not all genetic lesions are the actual drivers of tumor progression and determine therapy response^2,35,36^. Therefore, for this study we chose a well-defined system, the CRC tumor progression organoid model (TPO ^37^. The TPO model consists of organoids derived from healthy colon tissue that subsequently have been genetically engineered to harbor the four most frequent driver mutations of CRC (*APC^KO^, KRAS^G12D^, p53^KO^, SMAD4^KO^*). Here, we show that 5-FU toxicity relies on the active proliferation state of the cells, as it exerts its function by reducing TTP pool, which leads to DNA damage and cell death. In line with that, we observed that the actively proliferating CRC stem cell pool is efficiently targeted by 5-FU. Importantly, in this respect, we observed that p53 defines sensitivity to 5-FU: whereas p53-deficiency leads to DNA damage-induced cell death, active p53 protects from these effects through inducing G1 arrest. As we found that the 5-FU mode of action relies on a pyrimidine imbalance, we targeted the rewired metabolism of cancer cells to improve efficacy of the antimetabolite 5-FU. Of note, we found that rewiring glucose metabolism lowers the levels of nucleotides and enhances the efficacy of 5-FU in CRC cells without increasing toxicity in healthy non-transformed intestinal cells.

## Results

### 1. P53 determines sensitivity to 5-FU-induced DNA damage and cell death in human CRC organoids

To link genetic background and CRC progression state to 5-FU sensitivity, we analyzed response in organoid lines with defined, genetically engineered mutations: non-transformed (WT), *APC^KO^KRAS^G12D^* (AK), *APC^KO^P53^KO^* (AP), *APC^KO^KRAS^G12D^P53^KO^* (APK) and *APC^KO^KRAS^G12D^P53^KO^SMAD4^KO^* (APKS)^37^. This model has been studied *in vitro* and *in vivo* and APK and APKS organoids recapitulate morphological features of respectively adenocarcinoma and poorly differentiated adenocarcinoma with metastatic potential^37,38^. First, we analyzed sensitivity to 5-FU treatment and found that the response was dissimilar across the different organoid lines. 5-FU significantly reduced viability of AP, APK and APKS organoids. In contrast, WT and AK organoids showed no significant decrease in cell viability upon 5-FU treatment, although differences in organoid sizes were observed (Figure 1A and B, S1A). Interestingly, these results point at p53 having a protective role against 5-FU toxicity. P53 is a well-established factor and component of the DNA damage response and 5-FU can interfere with DNA synthesis^3,9,39^. Thus, we evaluated the impact of 5-FU on DNA damage by analyzing the DNA damage marker γH2AX. Western blot and flow cytometry analysis showed that, in line with the cell viability observations, 5-FU treatment induced DNA damage in the p53-deficient organoids, whereas this phenotype was milder and/or not significant in the WT and AK organoids (Figure 1C, D, S1A, B).

**Figure 1.**
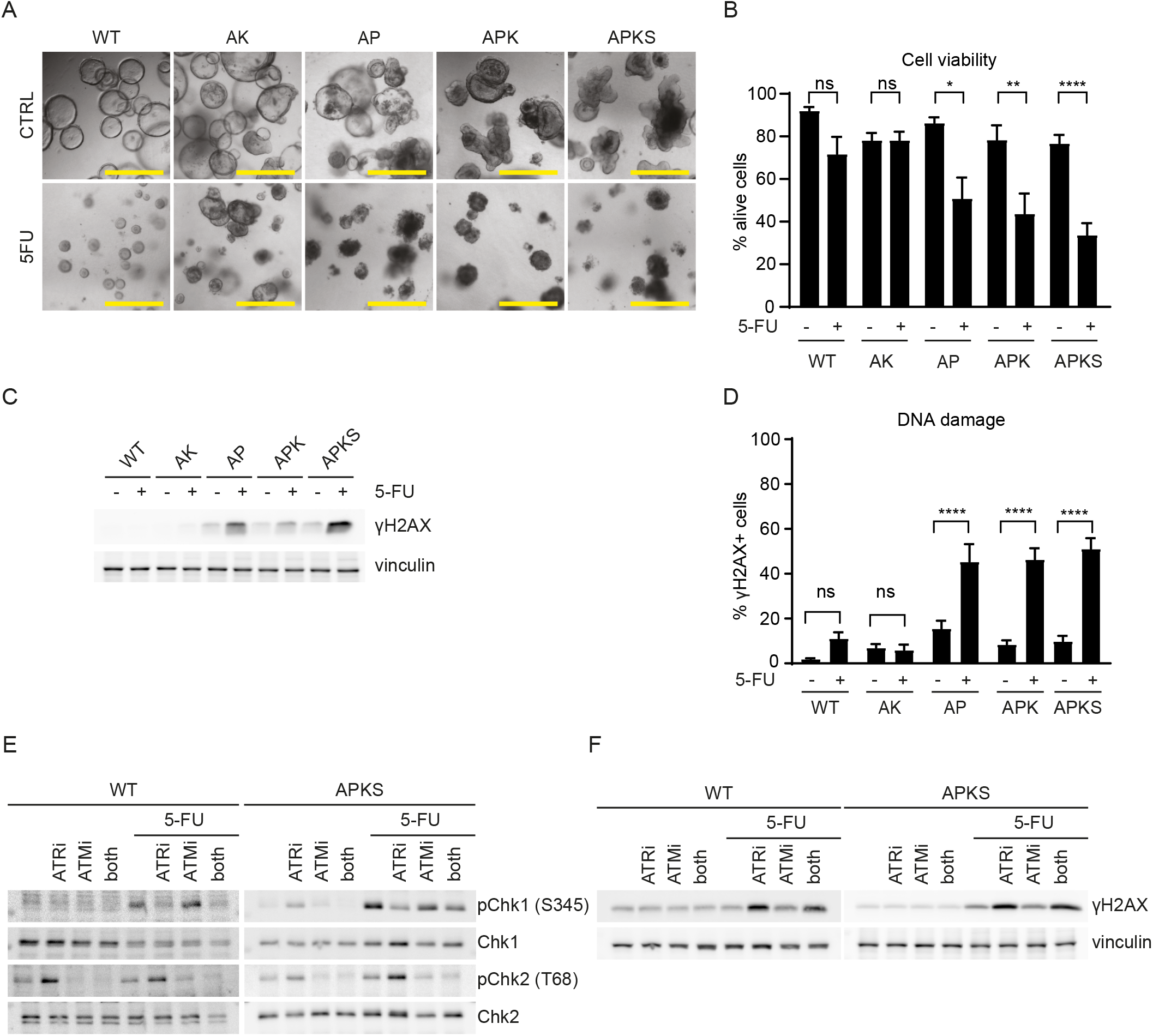
5-FU induces DNA damage and cell death in p53-deficient CRC organoids. **A)** Representative brightfield images of WT and CRC organoids treated with 5-FU for 7 days (scale bar = 500 μm). **B)** Cell viability analysis of WT and CRC tumor organoids by flow cytometry to distinguish alive (DAPI^-^) from death cells (DAPI^+^) upon 7 days of 5-FU treatment (mean ± SEM, n = 3-5, ANOVA, Sidak’s multiple comparisons test). **C)** Western blot detection of γH2AX and vinculin in lysates from WT and CRC tumor organoids treated with 5-FU for 48 hours (representative for n = 5). **D)** Quantification of cells with DNA damage by flow cytometry of WT and CRC organoids treated with 5-FU for 48 hours and stained with anti-γH2AX (mean ± SEM, n = 3-8, ANOVA, Sidak’s multiple comparisons test). **E,F)**, Western blot detection of (p)Chk1, (p)Chk2, γH2AX and vinculin of lysates from WT and APKS organoids treated with 5-FU for 24 hours and co-treated with either ATR inhibitor VE-821 or ATM inhibitor KU55933 or both for 26 hours (representative for n = 3, WT: Chk2: Santa Cruz, #SC-9064, APKS: Chk2: Cell Signaling, #3440). ns: non-significant, * p < 0.05, ** p < 0.01, **** p < 0.0001.

In order to gain further insights into the 5-FU-induced DNA damage, we analyzed activation of the different DNA damage response pathways. Stalled replication forks and single stranded DNA breaks are common consequences of conventional chemotherapies^40,41^. Hence, we first evaluated the ATR-Chk1 pathway, which is activated upon replication stress and single strand breaks^42^. Time course experiments revealed that 5-FU induces Chk1 activation in both WT and P53-deficient organoids as early as 24 hours. At 24 hours, WT organoids showed a mild induction of DNA damage marker γH2AX. P53 wild type organoids cleared the damage within the next 24 hours, while P53-deficient organoids showed accumulation of γH2AX at this later time point (Figure S1B). This suggests that while p53WT cells resolve 5-FU-induced DNA damage, p53-deficient organoids fail to do so. Interestingly, inhibition of ATR prevents Chk1 activation in both transformed and WT organoids, indicating that Chk1 is activated by 5-FU as a result of replication stress (Figure 1E)^43^. In line with that, inhibition of the ATR-Chk1 pathway enhances γH2AX levels in both WT and AKPS organoids (Figure 1F), showing that this pathway is essential for resolving DNA damage upon 5-FU treatment. During replication stress, unrepaired stalled replication forks can lead to double stranded DNA breaks and activation of the ATM-Chk2 pathway^42,44^. Hence, we analyzed Chk2 activation upon 5-FU. We found that 5-FU leads to ATM-dependent activation of Chk2 in APK and APKS, but not in WT, AK and AP organoids (Figure 1E, S1B)^45^. This supports the notion that high-grade tumors are less efficient in resolving replication stress and double stranded breaks. Taken together, these results show that ATR-Chk1 pathway is required for resolving 5-FU induced replication stress and importantly, that 5-FU treatment leads to increased levels of unresolved DNA damage and cell death in p53-deficient tumor organoids.

### 2. P53-deficient organoids undergo 5-FU-induced DNA damage and cell death due to the lack of G1 arrest

Although the importance of p53 in the 5-FU response has been investigated, a clear picture is yet to emerge^17,46–49^. While *in vitro* studies have shown that p53 is required for 5-FU-induced apoptosis^50–54^, other epidemiological studies show that p53 expression correlates with 5-FU resistance in stage III and IV patients^46,48,55^. However, p53 expression *per se* does not correlate with activity, especially when it is mutated. In our study, we find that p53 loss of function leads to 5-FU sensitivity, supporting the notion of a protective role of p53 against 5-FU. Therefore, we investigated the mechanism behind such phenotype. First, we examined whether p53 is activated upon 5-FU treatment. Western blot analysis showed that p53 and its transcriptional target p21 increase at 4 and 16 hours of 5-FU treatment in WT and AK organoids, respectively (Figure 2A). P53 is a key factor in response to cellular stress and can induce cell cycle arrest and apoptosis. Mechanistically, P53 regulates cell proliferation via its downstream target p21 that binds to and inhibits the cyclin/CDK complexes which phosphorylate Rb, thereby preventing its phosphorylation and consequent inhibition of G1/S transition (reviewed in^56^). Here, we found that 5-FU treatment enhances p21 expression and causes a complete loss of Rb phosphorylation in p53^*WT*^ organoids, whereas p53-deficient organoids show remaining Rb phosphorylation and lack of p21 induction (Figure 2B). In agreement, cell cycle profile analysis showed that 5-FU induces a G1 arrest in p53^*WT*^ organoids, whereas it causes S-phase accumulation in p53-deficient organoids (Figure 2C, S1A). Together with the reduced size upon 5-FU-treatment in WT and AK organoids (Figure 1A), the aforementioned points at p53-induced G1-arrest as the mechanism preventing 5-FU induced DNA damage. To test that, we substituted p53 function in p53-deficient organoids by arresting the cells in G1 by the CDK4/CDK6 inhibitor palbociclib^57^. Upon 24 hours of palbocilib treatment, most cells were arrested in G1 (Figure S2A). Hence, we started the 5-FU treatment after 24 hours of palbociclib. Interestingly, we found that palbociclib-induced G1 arrest in p53-deficient organoids completely prevented DNA damage upon 5-FU (Figure 2D). This indicates that cells acquire DNA damage during DNA replication. Moreover, palbociclib-induced cell cycle arrest completely rescued 5-FU-induced cell death (Figure 2E, F), indicating that DNA damage, rather than RNA toxicity, is the main contributor to 5-FU toxicity. Taken together, these results show that P53 protects against 5-FU through inhibition of cell cycle entry which prevents DNA damage and cell death.

**Figure 2.**
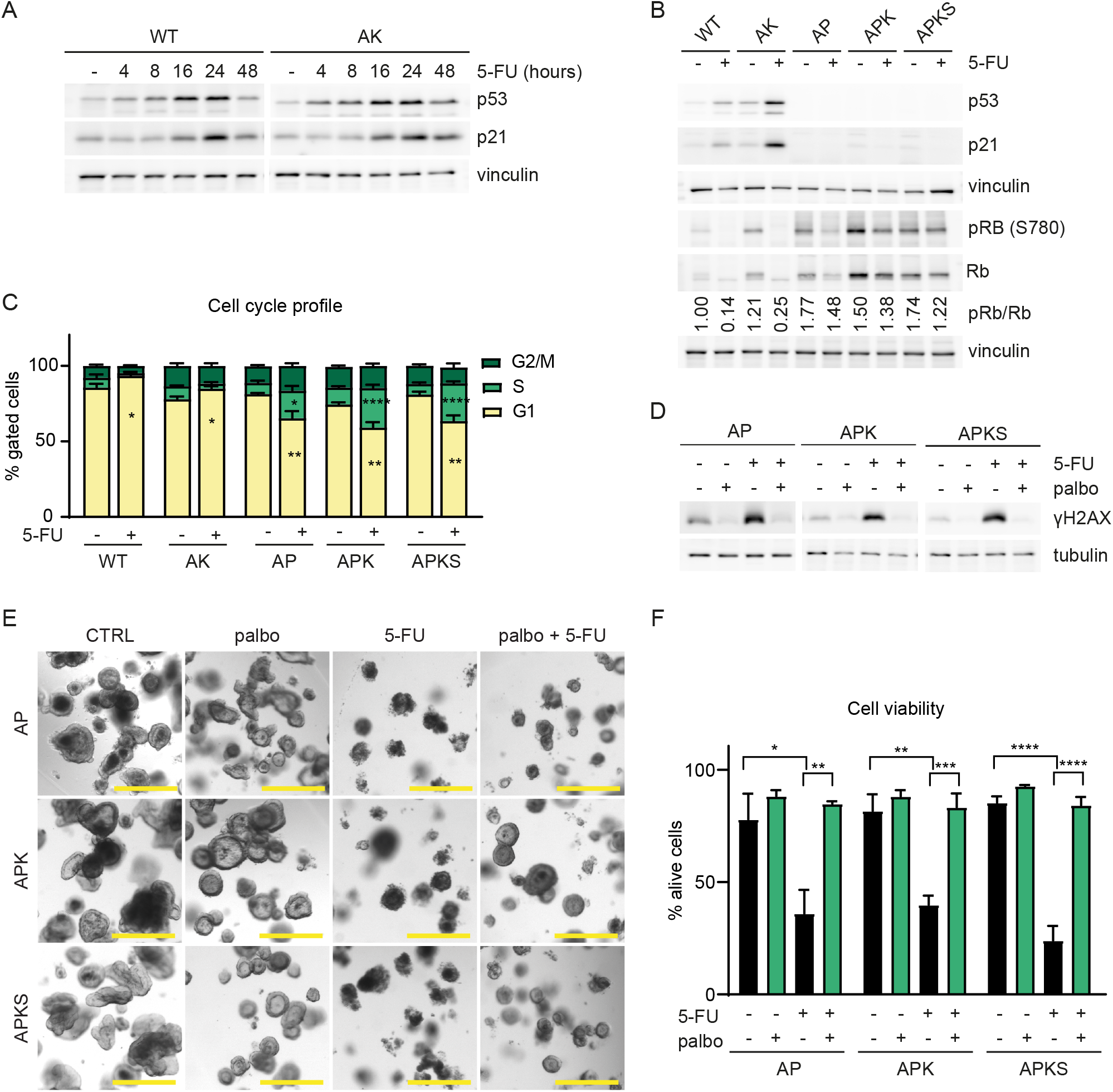
P53 protects against 5-FU-induced DNA damage and cell death via induction of a G1 arrest. **A)** Western blot detection of p53, p21 and vinculin in WT and AK organoids treated with 5-FU for 4, 8, 16, 24 and 48 hours (blot representative for n = 4). **B)** Western blot detection of p53, p21, (p)Rb and vinculin in WT and CRC organoids treated with 5-FU for 48 hours (representative for n = 4). **C)** Cell cycle profile determined by flow cytometry in WT and CRC organoids treated with 5-FU for 48 hours (mean ± SEM, n = 3-7, Kruskal-Wallis test, unpaired t-test and Mann Whitney test). **D)** Western blot analysis of γH2AX and tubulin of AP, APK and APKS organoids treated with 5-FU for 48hrs. Palbociclib treatment was started 24 hours before 5-FU treatment to arrest cells in G1 (representative for n = 3). **E, F)** Representative brightfield imaging (E) and cell viability analysis (F) of AP, APK and APKS organoids treated with 5-FU for 6 days. Palbociclib treatment was started 24 hours before 5-FU treatment to arrest cells in G1 (scale bar = 500 μm, mean ± SEM, n = 3, one-way ANOVA, Sidak’s multiple comparisons test). ns: non-significant, * p < 0.05, ** p < 0.01, *** p < 0.001, **** p < 0.0001.

### 3. 5-FU induces DNA damage in proliferating cancer (stem) cells

Our aforementioned results show that, although p53-deficient organoids are sensitive to 5-FU, not all cells respond to the treatment uniformly, as a fraction of cells do not show DNA damage and survive treatment (Figure 1B, D). To analyze the 5-FU response at a single cell level, we further investigated this on 5-FU responsive organoids (AP, APK and APKS). We first analyzed 5-FU-induced DNA damage by immunofluorescence and examined the response at the single cell level. Indeed, within single organoids, 5-FU-induced DNA damage is heterogeneous between the cells (Figure 3A, S3A). Next to genetic heterogeneity, CRC tumors display phenotypic heterogeneity as, similarly to the healthy intestine, the cells within the tumor show different degrees of stemness and proliferation rates^24,58^. In CRC tumors, CSCs are marked by high Wnt signaling, are mostly proliferative and fuel tumor growth^23–25,59^. To examine the response to 5-FU in CSCs, we genetically introduced in our organoid lines the Wnt-based stem cell reporter STAR^60–63^ (Figure 3B, S3B). Both in WT organoids and CRC organoids, cells showed heterogeneity in stemness (Figure 3B, S3B). Furthermore, EdU incorporation and cell cycle analyses showed that STAR^high^ cells in both in WT and CRC organoids, indeed constitute the proliferating population of cells (Figure S3C-E)^23,24^. Interestingly, flow cytometry and immunostaining analyses revealed that CSCs acquire more DNA damage than differentiated cells upon 5-FU treatment (Figure 3C, D and S3C, F), which relates to their high proliferation rate. In line with that, immunostaining of the proliferative marker Ki67 in combination with γh2ax, revealed that, upon 5-FU, KI67^pos^-proliferating cells have more DNA damage than non-proliferating Ki67^neg^ cells (Figure 3E, S3G). Furthermore, analysis of γh2ax in each of the cell cycle phases showed that the proportion of DNA damaged cells is higher in S and G2/M phase compared to cells in G1 (Figure 3F). Interestingly, this pattern observed in cycling CSCs, is also found in the (small proportion) of cycling STAR^low^, differentiated cells (Figure S3H). In line with G1-arrest having a protective effect against 5-FU, this result indicates that an active proliferation state in the cells is critical for 5-FU sensitivity.

**Figure 3.**
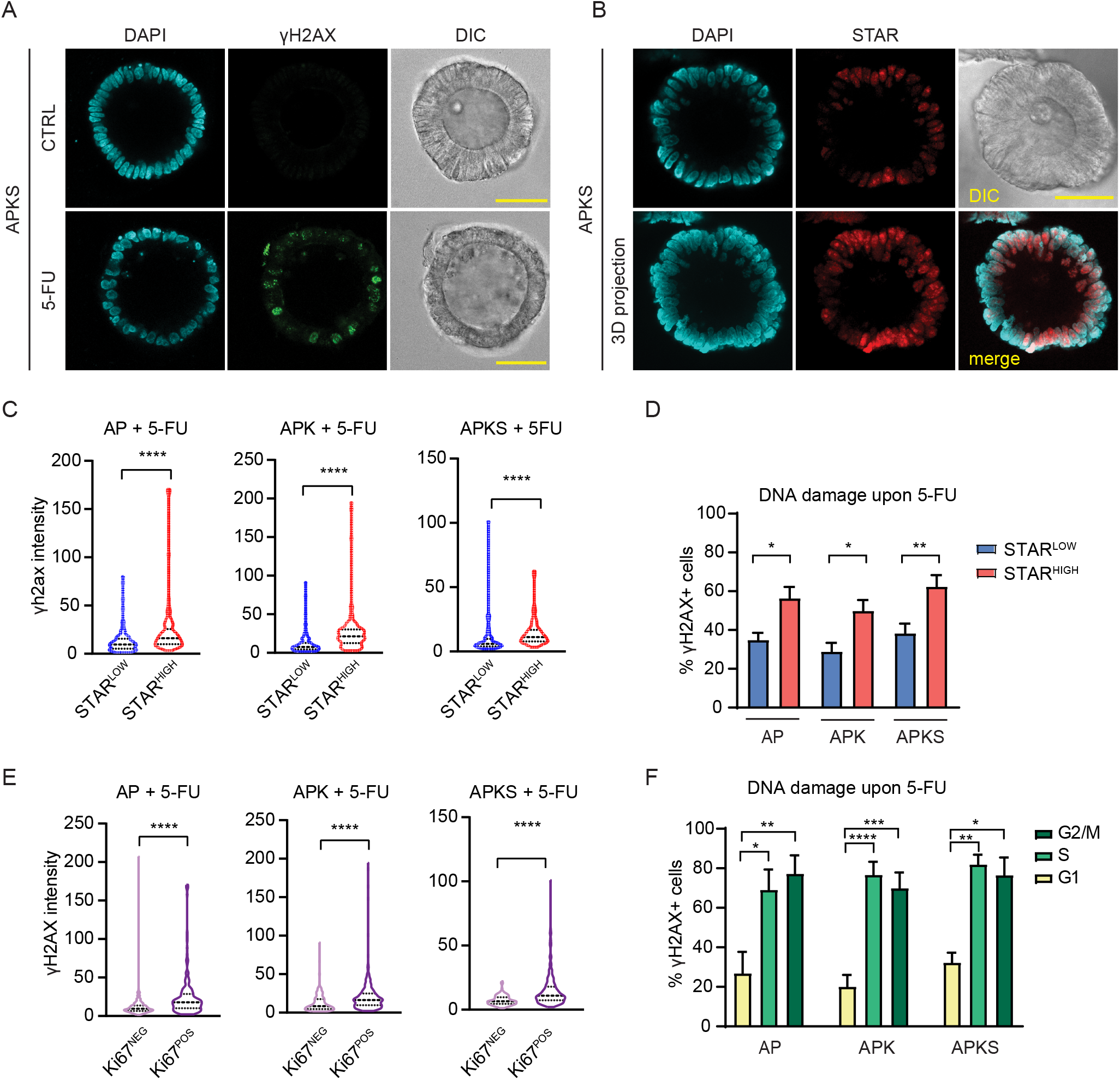
5-FU induces DNA damage in a proliferating subpopulation of tumor cells. **A)** Representative images of APKS organoids treated with 5-FU for 48 hours and stained with anti-γH2AX and DAPI (scale bar = 50 μm). **B)** Representative images of APKS organoids transduced with the stem cell reporter STAR and stained with DAPI (scale bar = 50 μm, upper panel: single Z-stack, lower panel: maximal projection of Z-stacks). **C)** Quantification of γH2AX intensity in STAR^low^ and STAR^high^ cells of immunofluorescent images of AP, APK and APKS organoids treated with 5-FU for 48 hours (Figure S3F) (70-153 cells per condition from 12 organoids from 3 independent experiments, Mann-Whitney test). **D)** Detection of γH2AX^+^ cells by flow cytometry in STAR^-^ vs STAR^+^ cells of AP, APK and APKS organoids treated with 5-FU for 48 hours (mean ± SEM, n = 6-7, one-way ANOVA, Sidak’s multiple comparisons test). **E)** Quantification of γH2AX intensity in KI67-positive vs KI67-negative cells of immunofluorescent images of AP, APK and APKS organoids treated with 5-FU for 48 hours (Figure S3G) (128-331 cells per condition from 12 organoids from 3 independent experiments, Mann-Whitney test). **F)** Detection of γH2AX^+^ cells by flow cytometry G1, S and G2/M cells of AP, APK and APKS organoids treated with 5-FU for 48 hours (mean ± SEM, n = 5-7, AP, APK: one-way ANOVA, Sidak’s multiple comparisons test, APKS: Kruskal-Wallis test, Dunn’s multiple comparisons test). ns: non-significant, * p < 0.05, ** p < 0.01, *** p < 0.001, **** p < 0.0001.

### 4. 5-FU induces DNA damage and cell death via a pyrimidine imbalance

The mechanism of 5-FU-induced cytotoxicity is still debated (reviewed in^3,9^). Here, we found that 5-FU-induced cell death relies on triggering DNA damage in cycling cells, which indicates replication stress. Based on the reported effect of 5-FU on TS activity, we analyzed whether 5-FU induced changes in the pyrimidine pool could explain this phenotype. First, we performed metabolomics analysis of APKS organoids, which showed the strongest response to 5-FU. We observed, that upon 5-FU, dUDP and dUMP levels increased, whereas the TDP and TTP pools were depleted (Figure 4A), indicating 5-FU inhibits TS activity in CRC organoids. Cancer cells have recently been described to exhibit a nucleotide overflow mechanism that can balance out disrupted pyrimidine synthesis^64^. This nucleotide overflow mechanism involves deoxyuridine excretion to prevent dUMP accumulation and the rate of deoxyuridine excretion is dependent on the extent of TS inhibition^64^. Interestingly, we observed an increase in extracellular deoxyuridine levels upon 5-FU treatment (Figure 4A), further supporting 5-FU induced TS inhibition. To examine the importance of the pyrimidine imbalance on DNA damage and cell death upon 5-FU, we performed nucleoside addback experiments. Addback of a mix of nucleosides (adenosine, guanosine, cytidine and thymidine) prevented 5-FU-induced DNA damage, S-phase accumulation and cell death in APKS organoids (Figure 4B, C, E and F). Interestingly, single addition of thymidine, but not the other nucleosides, rescued the 5-FU phenotype (Figure 4B, C). Thymidine administration by itself did not affect cell cycle progression, showing that this rescue is not dependent on any cell cycle effect (Figure 4D). Moreover, thymidine addback was sufficient to rescue 5-FU-induced cell death (Figure 4E, F). Together these results indicate that, mechanistically, 5-FU induces DNA damage and cell death by altering the pyrimidine pool.

**Figure 4.**
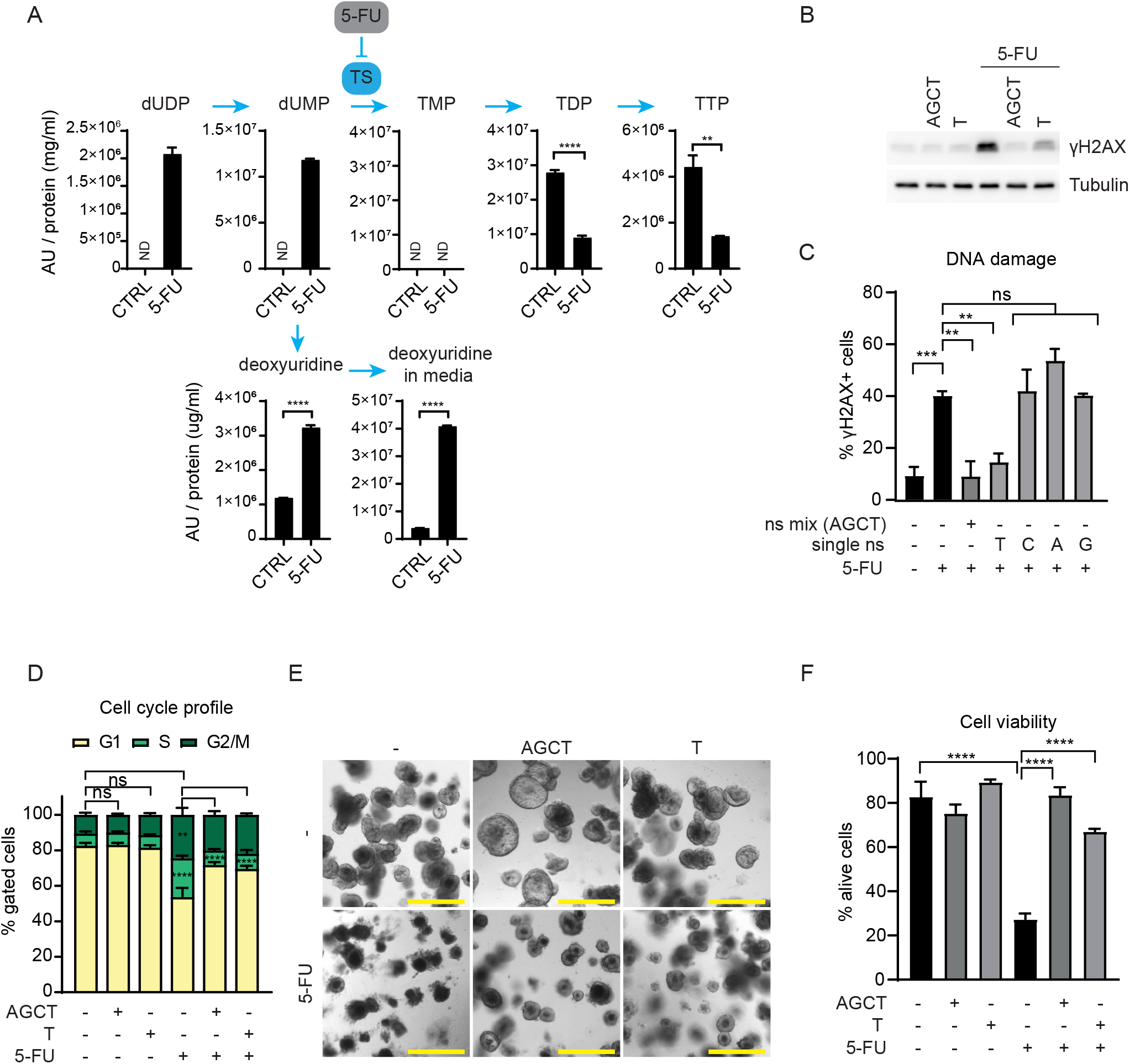
5-FU induces DNA damage and cell death by a pyrimidine imbalance. **A)** Detection of pyrimidines by metabolomics in APKS organoids treated with 5-FU for 30 hours (mean ± SEM, n = 3 technical replicates, one-way ANOVA, Sidak’s multiple comparisons test). **B)** Western blot detection of γH2AX and tubulin of APKS organoids treated with 5-FU, a nucleoside mix (adenosine, guanosine, cytosine and thymidine (25 μM each)), or thymidine (25 μM) for 48hrs (blot representative for n = 3)**. C)** Quantification of cells with DNA damage by flow cytometry of APKS organoids treated with 5-FU, a nucleoside mix (adenosine, guanosine, cytosine and thymidine (25 μM each)), or the separate nucleosides (25 μM) for 48hrs (mean ± SEM, n = 3-6, one-way ANOVA, Sidak’s multiple comparisons test). **D)** Cell cycle profile analysis by flow cytometry of APKS organoids, treated with 5-FU, a nucleoside mix (adenosine, guanosine, cytosine and thymidine (25 μM each)), or thymidine (25 μM) for 48 hours (mean ± SEM, n = 3, one-way ANOVA, Sidak’s multiple comparison’s test). **E, F)** Representative Brightfield images (E) and cell viability analysis (F) of APKS organoids treated with 5-FU, a nucleoside mix (adenosine, guanosine, cytosine and thymidine (25 μM each)), or thymidine (25 μM) for 6 days (scale bar = 500 μm, mean ± SEM, n = 3, one-way ANOVA, Sidak’s multiple comparisons test). Abbreviations: *A*: adenosine, *AU*: arbitrary unit, *C*: cytidine, *dUDP*: deoxyuridine diphospate, *dUMP*: deoxyuridine monophosphate, *G*: guanosine, *ND*: not detected, *ns mix*: nucleoside mix, ns: non-significant, *T*: thymidine, *TMP*: thymidine monophosphate, *TDP*: thymidine diphosphate, *TTP*: thymidine triphosphate, *TS*: Thymidylate synthase ** p < 0.01, *** p < 0.001, **** p < 0.0001.

### 5. Rewiring glucose metabolism lowers nucleotide levels and enhances the 5-FU-induced DNA damage

Derailed metabolism is one of the hallmarks of cancer. Cancer cells undergo the Warburg effect by avidly taking up of glucose and metabolize it through glycolysis into lactate independently of oxygen availability (Figure 5A). This increased glycolysis enables rapid production of ATP and supports the activity of anabolic pathways that rely on glycolytic intermediates, such as nucleotide synthesis. Based on the importance of nucleotides in the 5-FU toxicity, we rationalized that targeting the Warburg effect could improve 5-FU efficacy. First, we analyzed whether tumor organoids indeed undergo the Warburg effect by Seahorse bioenergetics analysis^65^. This showed that AK, APK and APKS organoids have higher glycolytic rates than non-transformed WT and AP organoids (Figure 5B), whereas the respiratory parameters are not significantly different (Figure S4A, B). These results show that the Warburg effect is recapitulated in *in vitro* organoids. Moreover, they point at constitutive active Ras signaling (*KRAS^G1C2^*) as the main driver of the Warburg effect, which is in line with previous studies (reviewed in^66^).

**Figure 5.**
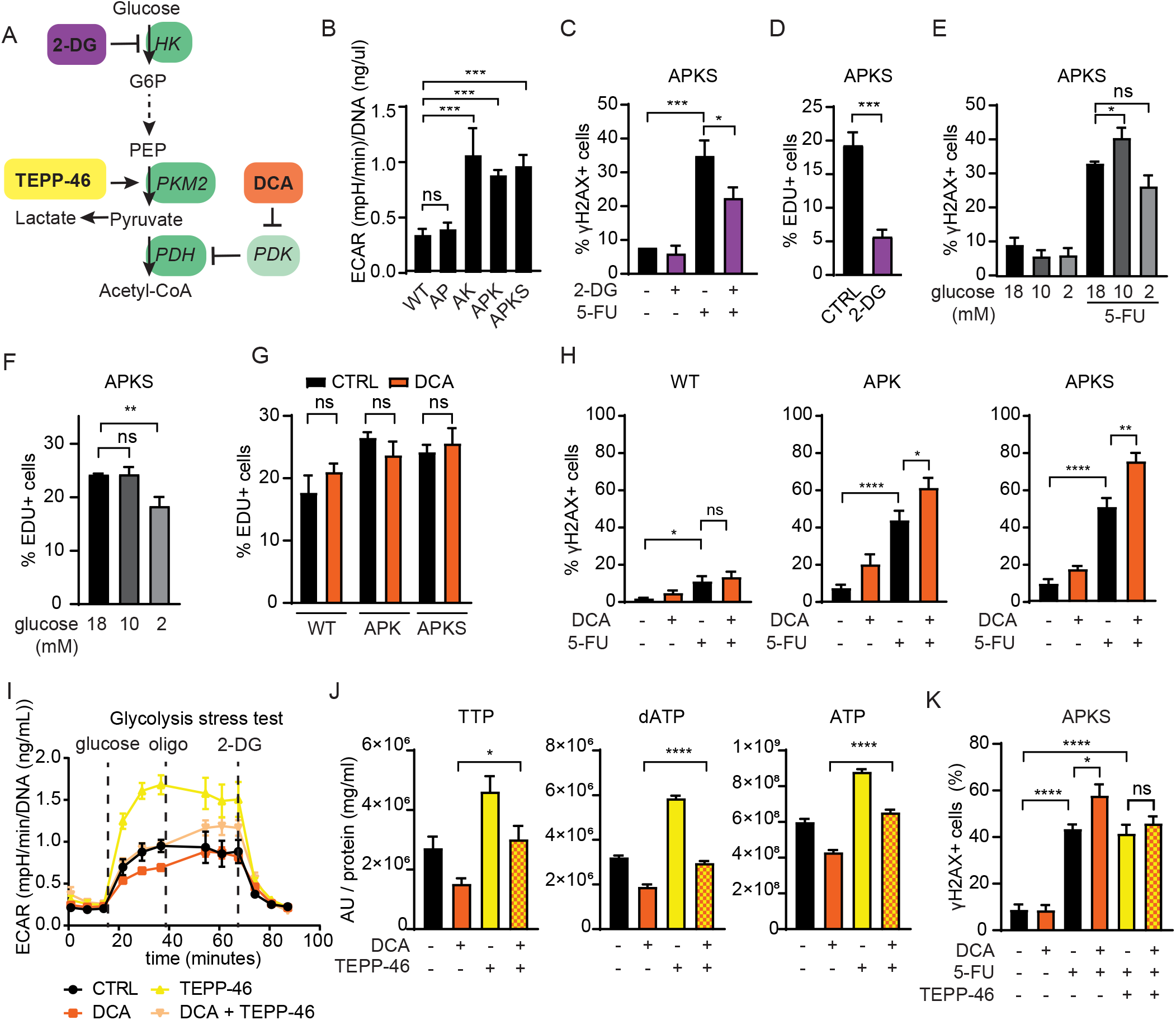
Rewiring glucose metabolism increases 5-FU-induced DNA damage. **A)** Schematic representation of glycolysis and different glycolysis-targeting drugs. **B)** Extracellular acidification rate (ECAR) of WT and CRC organoids determined by a glycolysis stress test by Seahorse XF analysis (mean ± SEM, n = 4-5, ANOVA, Sidak’s multiple comparisons test). **C)** Quantification of cells with DNA damage by flow cytometry of APKS organoids treated with 5-FU for 48 hours. 2-DG treatment started 20 hours before 5-FU treatment (mean ± SEM, n = 4, one-way ANOVA, Sidak’s multiple comparisons test). **D)** EdU incorporation analysis by flow cytometry of APKS organoids treated with 2-DG for 24 hours (mean ± SEM, n = 5, unpaired t-test). **E)** Quantification of cells with DNA damage by flow cytometry of APKS organoids treated with 5-FU for 48 hours. Glucose starvation started 20 hours before 5-FU treatment (mean ± SEM, n = 5-6, one-way ANOVA, Sidak’s multiple comparisons test). **F)** EdU incorporation analysis by flow cytometry of APKS organoids starved for glucose for 24 hours (mean ± SEM, n = 4, one-way ANOVA, Sidak’s multiple comparisons test). **G)** EdU incorporation analysis by flow cytometry of WT, APK and APKS organoids treated with DCA for 24 hours (mean ± SEM, n = 4-5, one-way ANOVA, Sidak’s multiple comparisons test). **H)** Quantification of cells with DNA damage by flow cytometry of WT, APK and APKS organoids treated with 5-FU for 48 hours. DCA treatment started 20 hours before 5-FU treatment (mean± SEM, n = 4-7, one-way ANOVA, Sidak’s multiple comparisons test). **I)** Determination of extracellular acidification rate (ECAR) during a Seahorse XF glycolysis stress test of APKS organoids treated with DCA and TEPP-46 for 24 hours (mean ± SEM, 5 technical replicates, representative for n = 3). **J)** Detection of TTP, dATP and ATP by metabolomics of APKS organoids treated with DCA and TEPP-46 for 24 hours (mean± SEM, 3 technical replicates, oneway ANOVA, Sidak’s multiple comparisons test). **K)** Quantification of cells with DNA damage by flow cytometry of APKS organoids treated with 5-FU for 48 hours. DCA and TEPP-46 treatments started 20 hours before 5-FU treatment (mean ± SEM, n= 3-4, one-way ANOVA, Sidak’s multiple comparisons test). Abbreviations: *2-DG*: 2-deoxyglucose, *ATP*: adenosine triphosphate, *dATP*: deoxyadenosine triphosphate, *DCA*: dichloroacetate, *G6P*: Glucose 6-phosphate, *HK*: Hexokinase, *PDH*: Pyruvate dehydrogenase, *PDK*: Pyruvate dehydrogenase kinase, *PEP*: Phosphoenol pyruvate, *PKM2*: Pyruvate kinase M2, *TTP*: thymidine triphosphate. ns: non-significant, * p < 0.05, ** p < 0.01, *** p < 0.001, **** p < 0.0001.

In order to lower the high glycolytic rates in tumor organoids, we first administered 2-deoxyglucose (2-DG), a glucose analogue that cannot be metabolized and accumulates in the cell leading to reduced glycolysis by the competitive inhibition of hexokinase-2 (HK2) (Figure 5A) (reviewed^67^). Although 2-DG efficiently decreased glycolysis (Figure S4C), in combination with 5-FU, it did not increase 5-FU-induced DNA damage and even lowered the effectiveness compared to 5-FU treatment on its own (Figure 5C). As 5-FU targets proliferating cells, we wondered whether 2-DG could alter proliferation. EdU incorporation analysis revealed that 2-DG indeed causes a major drop in proliferation (Figure 5D), suggesting that targeting glycolysis while affecting proliferation does not enhance the 5-FU effect. Next, we lowered glucose availability. Indeed, a limited reduction in glucose availability prior to 5-FU treatment increased 5-FU-induced DNA damage (Figure 5E). However, further lowering glucose concentration reduced cell proliferation and, consequently, 5-FU efficacy (Figure 5E, F). Precise regulation of glucose levels in the tumor microenvironment is rather challenging as a therapeutic strategy. Thus, we looked for pharmacological options to reduce glycolysis in tumor organoids. DCA (dichloroacetate) is a ‘low cost simple drug’, well tolerated and used in the treatment of acute and chronic lactic acidosis and metabolic diseases (reviewed in^68^). DCA is an inhibitor of pyruvate dehydrogenase kinases (PDKs), which increases pyruvate dehydrogenase (PDH) activity and consequently the conversion of pyruvate into acetyl-coA at the expense of lactate production, thus rewires glucose metabolism (Figure 5A). To test the effect of DCA in organoids, we analyzed PDH phosphorylation and bioenergetics by Seahorse analysis. DCA treatment decreased PDH phosphorylation in all organoid lines (Figure S4D), indicating that PDK activity was inhibited. Moreover, Seahorse analysis showed that DCA indeed reduces the high glycolytic rates of organoids that display the Warburg effect to glycolytic rates that are comparable to those of WT organoids (Figure S4E). Importantly, we analyzed the effect of DCA on proliferation and found no significant changes (Figure 5G). Next, we assessed whether DCA altered nucleotide metabolism by metabolomics and found that DCA treatment indeed lowered nucleotide levels (Figure 5J, S4F). Finally, as DCA lowered the glycolytic flux and the nucleotide levels without affecting proliferation, we expected on the basis of our results that DCA enhances the effect of 5-FU. Flow cytometry analysis revealed that, indeed, administration of DCA prior to 5-FU treatment increases the number of cells that undergo DNA damage upon 5-FU treatment in glycolytic APK and APKS organoids (Figure 5H, S4G).

Next, we evaluated whether the synergistic effect of DCA on 5-FU treatment indeed relies on the inhibition of glycolysis. Proliferating cancer cells exhibiting the Warburg effect often express the pyruvate kinase M2 isoform (PKM2)^69^. As PKM catalyzes the rate-limiting step of glycolysis, we evaluated whether TEPP-46, a specific activator of PKM2 could rescue the effects of DCA treatment (Figure 5A)^70^. To that end, we analyzed the glycolytic rates by Seahorse and the intracellular levels of glucose by live imaging of a glucose FRET sensor^71^. These analyses showed that PKM2 activation reverts glycolysis inhibition and restores intracellular glucose levels upon DCA treatment (Figure 5I, S4H-J). Importantly, in agreement with these results, PKM2 activation rescued the decrease in nucleotides resulting from DCA treatment (Figure 5J). Furthermore, flow cytometry analysis showed that, indeed, TEPP-46 treatment prevented the additive effect of DCA on 5-FU-induced DNA damage (Figure 5K). Altogether, this shows that DCA enhances the 5-FU induced DNA damage through lowering nucleotide levels as a consequence of inhibition of the Warburg effect.

### 6. Rewiring glucose metabolism enhances 5-FU-induced cell death

Next, we evaluated whether DCA in combination with 5-FU can further increase cell death. To that end, we treated WT and 5-FU responsive organoids (APK and APKS) with 5-FU and 5-FU/DCA. Qualitative analysis showed that while upon 5-FU treatment some organoids survive the treatment, those were absent in the DCA/5-FU treatment (Figure 6A). We quantitatively analyzed this phenotype in bulk by flow cytometry and at the single organoid level, by calculating a viability score based on imaging analysis. In both cases, we found that DCA in combination with 5-FU leads to reduced cell viability when compared to single 5-FU treatment (Figure 6B, C and S5A, B). Importantly, this additive effect of DCA was absent in WT organoids (Fig. 6A, C and S5A, B). Lastly, we evaluated the outgrowth capacity posterior to treatment, as a proxy for post-treatment survival and relapse potential. To that end, we treated organoids with 5-FU or with 5-FU/DCA, removed treatment to allow organoid recovery and re-plated them to analyze organoid formation capacity. Interestingly, we found that the 5-FU/DCA combination lead to a fewer number of formed organoids when compared to the 5-FU treatment in specifically in both APK and APKS organoids (Fig. 6D-E and S5C). Thus, targeting the Warburg effect in combination with 5-FU treatment increases DNA damage and cell death of highly glycolytic p53-deficient tumors without affecting non-transformed cells.

**Figure 6.**
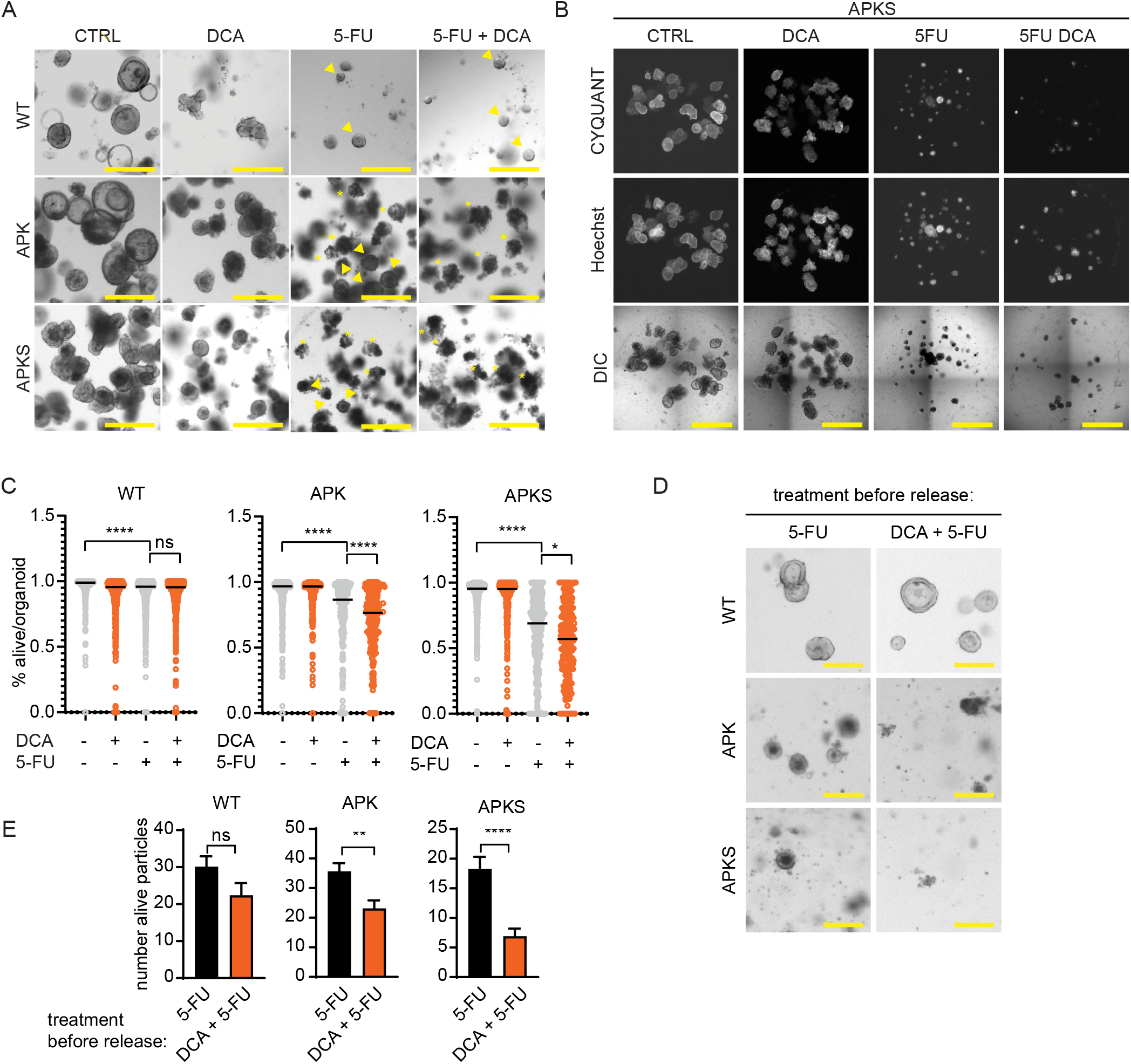
Rewiring glucose metabolism by DCA improves 5-FU-induced cytotoxicity. **A)** Representative brightfield images of WT, APK and APKS organoids treated with 5-FU for 7 days. DCA treatment was started 20 hours before 5-FU administration (scale bar = 500 μm, arrow heads indicate survivor organoids, asterisks indicate compromised organoids). **B)** Representative images of APKS organoids, stained with CYQUANT (alive) and Hoechst (total), treated with 5-FU for 7 days. DCA treatment started 20 hours before 5-FU treatment (scale bar = 1 mm). **C)** Quantification of alive score of images from B and S6B) (median, 237-720 organoids from 3 independent experiments, Kruskal-Wallis test, Dunn’s multiple comparisons test). **D)** Representative brightfield images of WT, APK and APKS organoids 4 days after replating, upon a 48 hour-5-FU treatment (50 μM) (with or without DCA treatment that started 20 hours before 5-FU administration), followed by 7 days of recovery time (scale bar = 200 μm). **E)** Determination of alive organoid particles based on S6C (mean ± SEM, WT: 8 matrigel droplets from 2 independent experiments, APK and APKS: 16 matrigel droplets from 4 independent experiments, WT and APK: unpaired t-test, APKS: Mann-Whitney test). ns: non-significant, * p < 0.05, ** p < 0.01, **** p < 0.0001.

## Discussion

Here we use a CRC tumor progression organoid model system to dissect the mode of action of 5-FU and to understand the importance of the different driver mutations on determining 5-FU efficacy in human colorectal tumors. We found that the efficacy of 5-FU is dependent on the proliferation status of cells and that it induces its toxicity via impairment of TTP synthesis. Interestingly, in this model, p53 has a protective role against 5-FU, as it inhibits G1/S transition and thereby prevents cells to acquire DNA damage during replication, and consequently prevents cell death. To enhance 5-FU efficacy, we targeted the Warburg effect with the purpose of further altering the nucleotide pool without affecting cell cycle progression. This strategy, as illustrated by the use of DCA, indeed improved 5-FU efficacy in p53-deficient glycolytic tumors, and importantly it shows no added toxicity to healthy non-transformed cells.

Precision medicine aims at patient stratification for treatment based on their tumor’s genetic background, in a way that cancer patients receive the most adequate treatment^72^. Stratification for targeted therapies has been facilitated by genomics. However, for conventional chemotherapies, such as 5-FU, this has been unsuccessful, partially due to the unresolved understanding of the mechanisms of action^2,19^. Accumulating evidence points at 5-FU inducing cytotoxicity via both DNA and RNA^3,9–14,73^. This apparent differential mode of action could arise from the dose of 5-FU (ranging from 1 to 1000 μM in these studies), where high 5-FU levels have been proposed to target cells through RNA toxicity, whereas prolonged exposure to low doses is proposed to be cytotoxic via TS inhibition-induced DNA damage^3,74–76^. In patients’ plasma and tumors, the (low) 5-FU concentrations (8 μM and 45 μM/kg respectively^77^), suggest that the DNA damaging effect more likely causes toxicity in CRC tumors. In line with that, TS expression and the 5-FU response do correlate, where tumors with high TS levels are commonly more resistant to 5-FU therapy than tumors with low TS levels^3,15–18^. In our CRC organoid system, we find that 5-FU induces cytotoxicity in cycling cells mainly via impaired pyrimidine synthesis-induced DNA damage accumulation. Although we have not observed 5-FU-induced cytotoxicity in non-cycling cells, different 5-FU concentrations or timings could reveal alternative mechanisms of 5-FU-induced cytotoxicity.

Identifying a role for specific driver mutations in drug efficacy is challenging^2,35,36^. Here, we find that p53 is a discriminating factor in 5-FU-induced toxicity, where p53-deficient organoids show sensitivity to 5-FU treatment. Previous studies have shown two scenarios for p53: a cell cycle arrest-dependent protective role against 5-FU^50,78,79^, but also that p53 is required to induce 5-FU-dependent apoptosis^14,50–54^. The balance between p53-induced cell cycle arrest and apoptosis could be dependent on 5-FU dose, the presence of additional mutations, cell intrinsic factors (epigenetic state, active signaling pathways) and cellular context (cellular microenvironment)^80–83^.

CSCs were defined as a subpopulation of cells within a tumor that have high tumor initiating capacity. Therefore, originally it was suggested that CSCs are responsible of resistance to therapy and tumor relapse and^21^. However, the occurrence of cellular plasticity by which non-CSCs can gain a CSC phenotype^24,59^, indicates that therapy resistance is unlikely to be attributed to a specific cell type. In CRC tumors, most CSCs show high Wnt signaling and active proliferation^23–25,59^. In this work, we find that CSCs are efficiently targeted by 5-FU which is due to their proliferative state. Interestingly, we observed a population of stem cells in G1 that do not accumulate DNA damage. This could be in line with the previously reported subpopulation of slow cycling/quiescent CSCs within the CSC subpopulation showing increased resistance and relapse potential in CRC tumors^84^. Together, these results stress that cellular behavior rather than being a specific cell type determines 5-FU-sensitivity^81^.

In cancer the metabolism of the cells is rewired to support their uncontrolled proliferation^85^. This rewiring can be caused by different mutations, but always serves the goal to support proliferation (reviewed in^85–88^). The Warburg effect has been recently revised and it is proposed that increased glycolysis with lactate as end product balances the cellular redox state to favor anabolic pathways (reviewed in^85,86^). DCA is an FDA-approved drug that is already used for the treatment of some metabolic diseases including lactic acidosis with minimal side effects^68,89^. Here, we show that DCA targets the Warburg effect by reducing glycolysis and lowering the nucleotide levels in glycolytic CRC tumors, while proliferation is maintained, which is key for enhancing 5-FU efficacy. We found that administration of 5-FU upon DCA-induced metabolic rewiring improved its efficacy APK and APKS organoids, which form respectively differentiated and poorly differentiated adenocarcinomas *in vivo*^38^. Based on its mode of action, we propose that DCA can also improve the efficacy of other replication-targeting chemotherapies. Interestingly, while our results show that DCA improves efficacy in 5-FU sensitive cells, former studies showed that DCA could restore 5-FU sensitivity to cells with acquired resistance^90,91^.

While precision medicine moves forward, conventional chemotherapies are still the workhorse of oncology. A better understanding of their mode of action is key to find ‘low cost-low toxicity’ strategies to improve patient treatment. Considering that DCA is a low cost drug and does not have adverse effects in healthy tissue, it emerges as a promising strategy of improving conventional chemotherapies.

## Materials and Methods

### Organoid culture

Tumor progression organoids were a gift from the Drost lab ^37^. Organoids were cultured at 37°C and at 5% CO2. A mycoplasma-free status was confirmed routinely. The basic culture medium contained advanced DMEM/F12 supplemented with penicillin/streptomycin, 10 mM HEPES and 20 mM Glutamax. For experiments, upon trypsinization into single cells/small clumps of cells, cells were cultured in matrigel (Corning, #356231) or BME (Bio-Techne, #3533-010-02) in expansion medium (Table 1), supplemented with Rock inhibitor Y-27632 (Gentaur, #607-A3008).

**Table 1.** Expansion medium composition

|  | WT | AK | AP | APK | APKS |
| --- | --- | --- | --- | --- | --- |
| 0.25 nM Wnt surrogate FC fusion protein (U-protein express BV, #N001) | x |  |  |  |  |
| 20% v/v R-spondin1-CM (homemade) | x |  |  |  |  |
| 10% v/v Noggin-CM (homemade) | x | x | x | x |  |
| 1x B27 (Fisher Scientific, #15360284) | x | x | x | x | x |
| 10 mM Nicotinamide (Sigma, A9165) | x | x | x | x | x |
| 1.25 mM N-acetyl-cysteine (Sigma, #N0636) | x | x | x | x | x |
| 3 μM SB202190 (Gentaur, #607-A1632) | x | x | x | x | x |
| 500 nM A83 (Biotechne, #2939) | x | x | x | x | x |
| 50 ng/mL EGF (Peprotech, #AF-100-15) | x |  | x |  |  |

#### Experimental set-up

After 7 days, organoids were diluted and replated in 24-well plates (4 droplets of 11 μl) and medium was replaced to differentiation medium (Table 2). After 3 days, differentiation medium was refreshed and after another 20 hours, 5-FU (Sigma, #F6627) was added. For all experiments 100 μM of 5-FU was used, unless stated differently. The timing of the 5-FU treatments is stated in the figure legends.

**Table 2.**
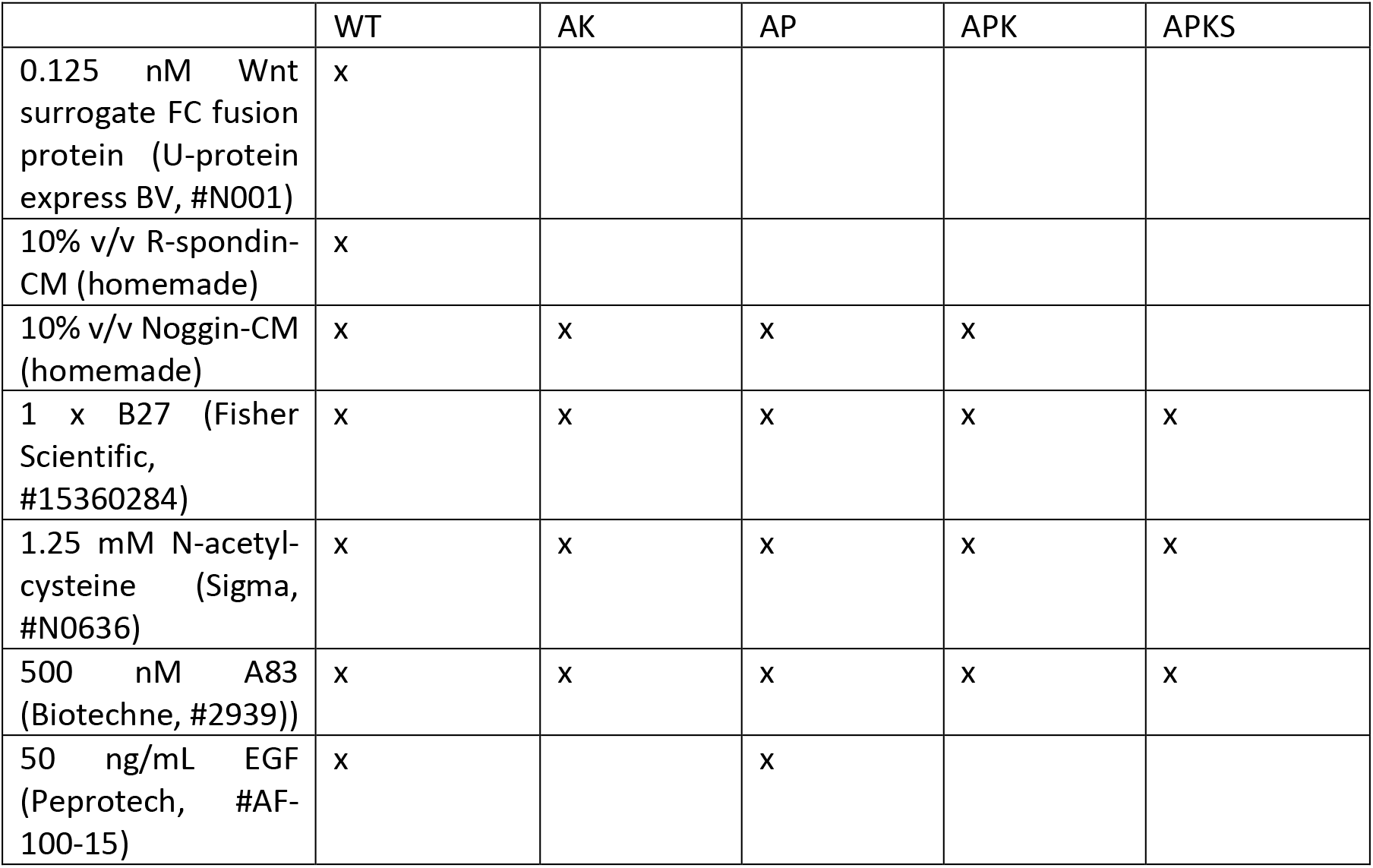
Differentiation medium composition

Treatments with ATR inhibitor VE-821 (5 μM, Bioconnect, #S8007) and ATM inhibitor KU-55933 (10 μM, Sigma, #SML1109) were started 2 hours before 5-FU administration. Nucleosides (25 μM each, all Sigma: adenosine #A4036, thymidine #T9250, guanine #G6264 and cytidine #C4654) were added together with 5-FU. Treatments with palbociclib (1 μM, selleckchem, #S1116)) were started 24 hours before 5-FU addition. DCA (20 mM, Sigma, #347795), 2-DG (100 mM, Sigma #D8375) and TEPP-46 (100 μM, Selleckchem, #S7302 treatments) or glucose starvations were started 20 hours before 5-FU treatment. Glucose starvation medium was prepared with SILAC Advanced DMEM/F-12 Flex Media (Gibco, #A2494301), supplemented penicillin/streptomycin, 10 mM HEPES and 20 mM Glutamax, L-Arginine (147,5 mg/L, Sigma, #A6969), L-Lysine (91,25 mg/L, Sigma, L8662) and the stated glucose (Merck Millipore, #1.08337.1000) concentration. For EdU incorporation analysis, 6 hours prior to collection organoids were incubated with 1 μM EdU (Thermo fisher, #C10636).

### Lentiviral transduction

From the Stem cell ASCL2 reporter (STAR) plasmid ^60,62^, the 8x STAR-sTomato-NLS sequence was cloned into a lentiviral vector with a puromycin-resistance cassette. The pcDNA3.1 FLII12Pglu-700uDelta6 (Addgene plasmid #17866) was a gift from Wolf Frommer ^71^. The eCFP sequence was replaced for a codon optimized eCFP sequence to prevent recombination with the YFP sequence. The new glucose sensor sequence was cloned into a lentiviral vector under the control of a Hef1 promoter and with a puromycin resistance cassette. These constructs, together with third generation packaging vectors, HEK293T cells and LentiX Concentrator (Clontech) were used to generate and concentrate lentiviral particles. Organoids were lentivirally transduced as described in ^92^). In brief, organoids were trypsinized and incubated with concentrated virus (60 min while centrifuging at 600 rpm at RT followed by 4 h at 37°C). Next, organoids were plated in matrigel.

### Protein lysates & western blot

Organoids were washed once with and collected in ice-cold PBS, supplemented with 5 mM NaF (Vwr, #1.06449.0350) and 1 mM NaVO_3_ (Sigma, #S6508). Organoids were centrifuged and pellets were incubated in Cell Recovery Solution, supplemented with NaF and NaVO_3_ on ice for 15 minutes. Upon centrifugation, pellets were lysed in lysis buffer (50mM Tris pH 7.0, 1%TX-100, 15 μM MgCl_2_, 5μM EDTA, 0.1 mM NaCl, 5 mM NaF, 1 mM NaVO_3_, 1 μg/mL Leupeptin (Sigma, #11034626001) and 1 μg/mL Aprotinin (Sigma, #10981532001) and protein content was determined by Biorad protein assay (#500-0006). Samples were adjusted for the protein content and Laemli sample buffer was added. Proteins were run in SDS-PAGE and transferred to Immobilon Polyscreen PVDF transfer membranes (#IPVH00010, Merck Millipore) or Amersham Protan nitrocellulose membranes (#10600001 GE Healthcare Life Sciences). Western blot analysis was performed with primary antibodies recognizing: vinculin (Sigma, #V9131), Tubulin (Merck Millipore, #CP06 OS), γH2AX (Sigma, #05-636), pChk1(S345) (Cell Signaling, #2348, Chk1 (Santa Cruz, #SC-8408), pChk2(T68) (Cell Signaling, #2661), Chk2 (Cell Signaling, #3440 and Santa Cruz, #SC-9064), pRb(S780) (Cell Signaling, #9307), Rb (Santa Cruz, #SC-7905), p53 (Santa Cruz, #SC-126), p21 (BD Bioscience, #556430), pPDH(S293) (Abcam, #ab92696), PDH (Invitrogen, #459400). Secondary HRP-conjugated antibodies targeting mouse and rabbit IgG were purchased from Biorad.

### Flow cytometry

Organoids were collected in ice-cold DMEM/F12 medium, supplemented with penicillin/streptomycin, 10 mM HEPES and 1x Glutamax (DMEM/F12 +++ medium) and subsequently incubated with trypsin (Sigma). For cell viability analysis, cells were stained with DAPI (Sigma-Aldrich, #D9564) on ice for 5 minutes and immediately analyzed by flow cytometry. Viability was determined by DAPI staining, where DAPI^-^ cells were considered alive, and DAPI^+^ cell s as dead.

For γH2AX staining, cell cycle profile analysis and EdU incorporation analysis, single cells were fixed in (PFA) (#1004965000, Merck Millipore) at RT for 10 min and permeabilized overnight on ice with 70% EtOH. For γH2AX staining, cell cycle profile analysis, organoids were washed once with 10 mL PBS + 1% BSA and 0.02% tween, incubated with Phospho-Histone H2A.X (Ser 139)-Alexa Fluor 488 (eBioscience, #53-9865-82) for 30 min at RT, covered from light. For cell cycle profiling, organoids were incubated with RNAse (100 μg/ml) in PBS for 20 minutes at RT and with DAPI for 2 hours, on ice, covered from light. For EdU detection, cells were stained by the Click-iT Plus EdU Pacific Blue Flow cytometry assay kit (Thermo fisher, #C10636) according to manufacturer’s instructions. Flow cytometry was performed by using a BD FACS Celesta #660345.

### Immunofluorescence and life imaging

Organoids were washed once in ice-cold-PBS, were collected in 1 mL ice-cold cell recovery solution (Corning, #354253) and 1 mL of ice-cold PBS in a 15 mL tube, and were incubated on ice for 10 minutes. Organoids were washed once with ice-cold PBS and were fixed by 4% PFA (#1004965000, Merck Millipore) for 20 min at RT and stored in PBS at 4°C. For staining, organoids were transferred into 1.5 mL Eppendorf tubes. Organoids were permeabilized with PBS buffer containing 10% DMSO, 2% Triton X-100 and 10 g l^−1^ BSA for 4 hours at 4°C. Organoids were stained with primary antibodies (γH2AX 1:400, Sigma, #05-636, and Ki67 1:200, Abcam, #ab15580) overnight, Alexa fluorophore-conjugated secondary antibodies (Invitrogen) for 4 hours and with DAPI for 1 hour at 4 °C. Imaging was performed using a SP8 confocal microscope (Leica Microsystems). Light microscopy was performed using EVOS M5000 imaging system (Invitrogen).

#### Image analysis immunofluorescence

Image analysis was performed in ImageJ. For image analysis, images were converted into 32-bit images. Nuclei were automatically detected by the Stardist-2D macro based on the DAPI staining. Within these ROIs, intensities of gH2AX, KI67 or STAR were determined. STAR^low^ and STAR^high^ cells were identified as the nuclei with respectively 20% lowest and highest STAR intensities per organoid. For Ki67 positivity, a cutoff of intensity 20 was used.

#### Image analysis glucose sensor

Imaging of the FLII12Pglu-700uDelta6 glucose FRET sensor was performed in 4 independent experiments. Data analysis was performed in ImageJ. Images of YFP and CFP channels were converted into 32-bit images, automatic thresholding was performed and YFP/CFP ratio was visualized by using the image calculator tool. YFP/YFP ratios were quantified by measuring the mean of the ratios in whole organoids.

### Cell viability imaging

7 days after trypsinization, organoids were replated into a 96-well plate (5 μL matrigel/well, 3 wells/condition). For cell viability analysis, organoid were stained with CyQUANT Direct assay (Thermofisher, #C35011) according to the manufacturer’s instructions to detect the alive cells and Hoechst (#H1399, Life Technologies) to detect all dead and alive cells, for 1 hour at 37 °C. Organoids were imaged at a Cell observer Z1 (Zeiss).

#### Image analysis

In imageJ, images were converted into 32-bit, and maximal projections of C1 (CyQUANT) and C3 (Hoechst) images were created. Organoids in both channels were automatically detected by the Stardist-2D macro to generate regions of interest (ROIs) reflecting respectively the alive parts (alive ROIs) and the total organoids (total ROIs). In the C1 images, the alive ROIs were masked and converted into binary images. Now, in the binary C1 images, the total ROIs were imported, and the %area of alive ROIs was determined within these total ROIs and used as a viability score per organoid.

### Outgrowth experiments & imaging

7 days after trypsinization, organoids were replated into a 24-well plate (4 droplets of 11 μL) and cultured on differentiation medium. Upon 3 days, medium was refreshed and DCA (20mM) was added. After 20 hours, 5-FU (50 μM) was administrated and organoids were cultured for 48 hours. Then drugs were washed away and organoids were cultured for another 7 days on differentiation medium. After 7 days, organoids were trypsinized into small clumps and replated in matrigel in a 24-well plate and in a 96-well plate (5 μL matrigel/well, 4 wells/condition). Upon 4 days, organoids in 24-well plate were imaged by EVOS. Organoids in 96-well plate were stained with CyQUANT Direct assay (Thermofisher, #C35011) according to the manufacturer’s instructions and Hoechst for 1 hour at 37 °C. Organoids were imaged at a Cell observer Z1 (Zeiss).

#### Image analysis

In imageJ, images were converted into 32-bit, and maximal projections of C1 (CyQUANT) images were created. Alive organoid (particles) were automatically detected by the Stardist-2D macro.

### Seahorse XF Flux analysis

Seahorse Bioscience XFe24 Analyzer was used to measure extracellular acidification rates (ECAR) in milli pH (mpH) per min and oxygen consumption rates (OCR) in pmol O2 per min as previously described ^65^. In short, organoids were seeded in 3 μL matrigel per well in XF24 cell culture microplates (Seahorse Bioscience). 1 hour before the measurements, culture medium was replaced and the organoids were incubated for 60 min at 37°C. For the mitochondrial stress test, culture medium was replaced by Seahorse XF Base medium (Seahorse Bioscience), supplemented with 10 mM glucose (Sigma-Aldrich), 2 mM L-glutamine (Sigma-Aldrich), 5 mM pyruvate (Sigma-Aldrich) and 0.56 μL ml-1 NaOH (1M). During the test, 5 μM oligomycin, 2μM FCCP and 1μM of rotenone and antimycin A (all Sigma-Aldrich) were injected to each well after 18, 45 and 63 min, respectively. For the glycolysis stress test, culture medium was replaced by Seahorse XF Base medium, supplemented with 2 mM L-glutamine and 0.52 μL mL^−1^ NaOH (1 M). During the test 10 mM glucose, 5 μM oligomycin and 100 mM 2-deoxyglucose (Sigma-Aldrich) were injected to each well after 18, 36 and 65 min, respectively. After injections, measurements of 2 min were performed in triplo, preceded by 4 min of mixture time. The first measurements after oligomycin injections were preceded by 5 min mixture time, followed by 8 min waiting time for the mitochondrial stress test and 5 min mixture time followed by 10 min waiting time for the glycolysis stress test. OCR and ECAR values per group were normalized to the total amount of DNA present in all wells of the according group.

### Metabolomics

#### Materials

Organic solvents were ULC-MS grade and purchased from Biosolve (Valkenswaard, The Netherlands). Chemicals and standards were analytical grade and purchased from Sigma-Aldrich (Zwijndrecht, The Netherlands). Water was obtained on the day of use from a Milli Q instrument (Merck Millipore, Amsterdam, The Netherlands).

#### Sample preparation and LC-MS analysis

For metabolomics, 3 wells with 200 μL of organoid-containing matrigel were used per condition. During treatment, medium & drugs were refreshed 7 hours before collection. Upon collection time, from each well, 500 μl medium per well was collected, pooled with medium from the other wells from the same condition and snap-frozen in liquid nitrogen in 2mL Eppendorf tubes. Organoids from the same condition were pooled during collection. Organoids were washed once with ice-cold PBS, and subsequently collected in ice-cold PBS in a 15 mL falcon tube. Upon another wash with ice-cold PBS, organoids were transferred to 2 mL Eppendorf tubes, centrifuged, resuspended in 80% ice-cold methanol and snap-frozen in liquid nitrogen.

Samples were evaporated to dryness in a Labconco Centrivap (VWR, Amsterdam, The Netherlands). To the residue 350 μL water, 10 μL 1 mM ribitol internal standard in water, 375 μL methanol and 750 μL chloroform were added. After pulse vortex mixing, the samples were incubated for 2 hours in a VWR thermostated shaker (900 rpm, 37°). After centrifugation at room temperature (10 min, 15000xg) the upper aqueous phase was quantitatively transferred to a clean 1.5 μL Eppendorf tube and evaporated to dryness overnight in the Labconco Centrivap. The residue was dissolved in 100 μL water, transferred to an injection vial and kept at 6 °C during LC-MS analysis.

The LC-MS analysis was performed using a 2.1×100 mm Atlantis premier BEH-C18 AX column (2.1×100, 2.5 μm) connected to a VanGuard column, both purchased from Waters (Etten-Leur, The Netherlands). The column setup was installed into an Ultimate 3000 LC system (Thermo Scientific, Breda, The Netherlands). The column outlet was coupled to a Thermo Scientific Q-Exactive FT mass spectrometer equipped with an HESI ion source. The UPLC system was operated at a flow rate of 250 μL min^−1^ and the column was kept at 30 °C. The mobile phases consisted of 10 mM ammonium acetate and 0.04(v/v) ammonium hydroxide in water, pH9 (A), and acetonitrile (B), respectively. Upon 5 μL sample injection the system was kept at 0% B for 1 min followed by a 4 min linear gradient of 0-30% B. Thereafter, the gradient increased linearly to 95% B in 3 min and kept at 95% for 2 min. The column was regenerated at 0% B for 6 min prior to a next injection. All samples were injected 3 times. Mass spectrometry data were acquired over a scan range of m/z 72 to 900. The system was operated at −2.5 kV (negative mode) and 120000 mass resolution. Further source settings were: transfer tube and vaporizer temperature 350°C and 300°C, and sheath gas and auxiliary gas pressure at 35 and 10, respectively. For high mass accuracy mass calibration was performed before each experiment. Raw data files were processed and analyzed using XCalibur Quan software.

### Statistical Analysis

Statistical analysis for image analysis, flow cytometry and metabolomics results was performed by using Graphpad Prism 8. First Gaussian distribution of data was tested by a Shapiro-Wilk test to next apply parametric or non-parametric statistics. Details of statistics are described in the figure legends.

## Author contributions

Conceptualization, M.J.R.C., M.C.L., and B.M.T.B.; Methodology and Investigation, M.J.R.C., M.C.L., M.M., S.G., N.T.B.N.; and S.K.S.R; Metabolomics: M.C.G and E.C.A.S; Development tumor progression organoid model: J.D. and H.C.; Manuscript Writing, M.J.R.C. and M.C.L.; Review & Editing, M.J.R.C. and B.M.T.B.; Funding Acquisition, M.J.R.C. and B.M.T.B.

## Acknowledgements

We thank H.J.G. Snippert (UMC Utrecht) for sharing the stem cell activity reporter plasmid; W. Frommer (Heinrich-Heine-University) for sharing the glucose FRET sensor plasmid; I. Verlaan (UMC Utrecht) for preparing R-spondin- and Noggin-conditioned medium; and A. Janssen and J. Lehman for critical reading of the manuscript. This work was financially supported by Dutch Cancer Society (KWF 2016-I 10471, KWF 2017-II 11315).

---

**Figure S1.**
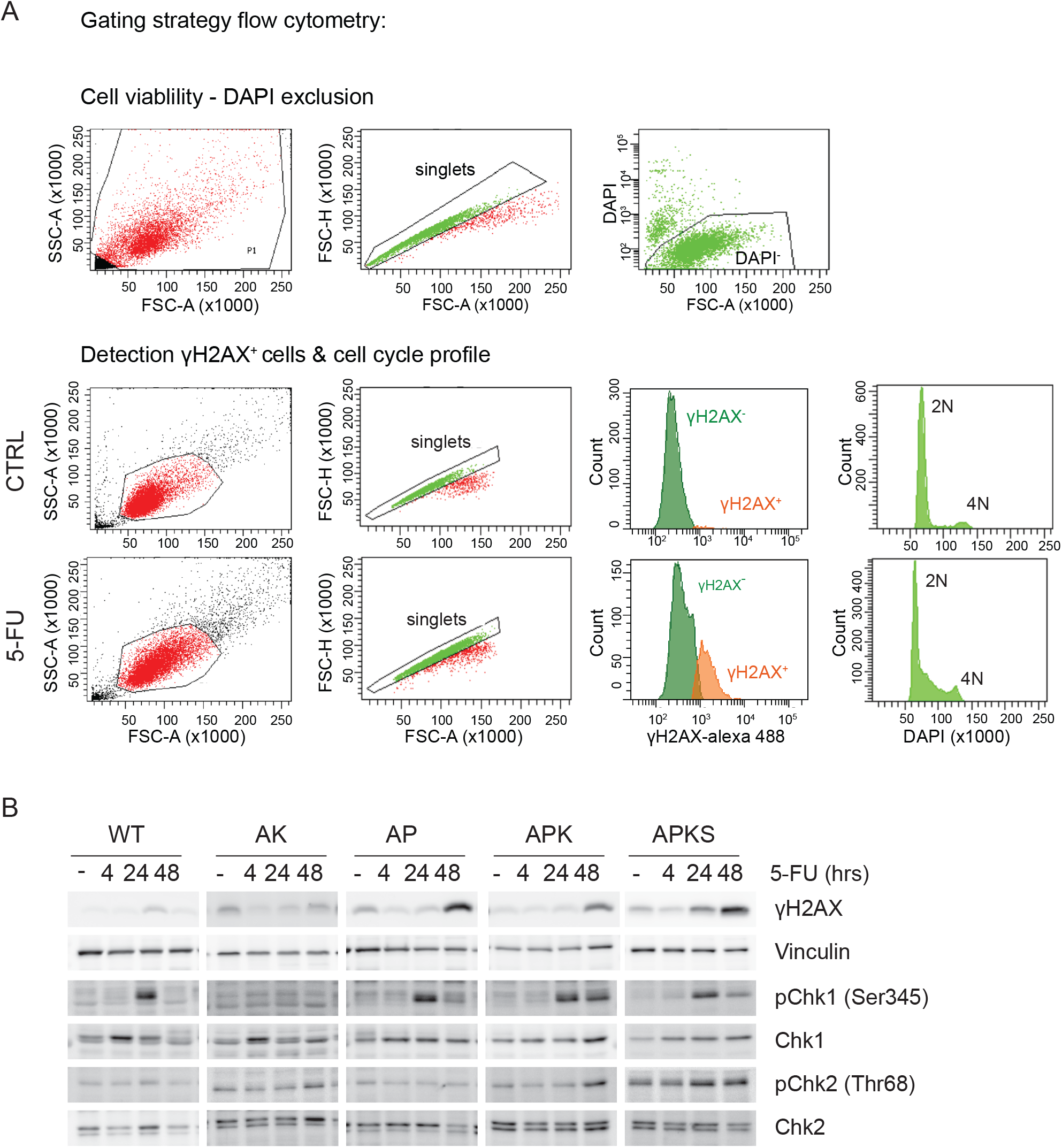
Flow cytometry gating strategy & 5-FU time course. **A)** Flow cytometry gating strategies for cell viability analysis by DAPI exclusion, detection of γH2AX^+^ cells and cell cycle profile analysis. **B)** Western blot detection of (p)Chk1, (p)Chk2, γH2AX and vinculin of lysates from WT and CRC organoids treated with 5-FU for 4, 24 or 48 hours (representative for n = 3, Chk2: Santa Cruz, #SC-9064).

**Figure S2.**
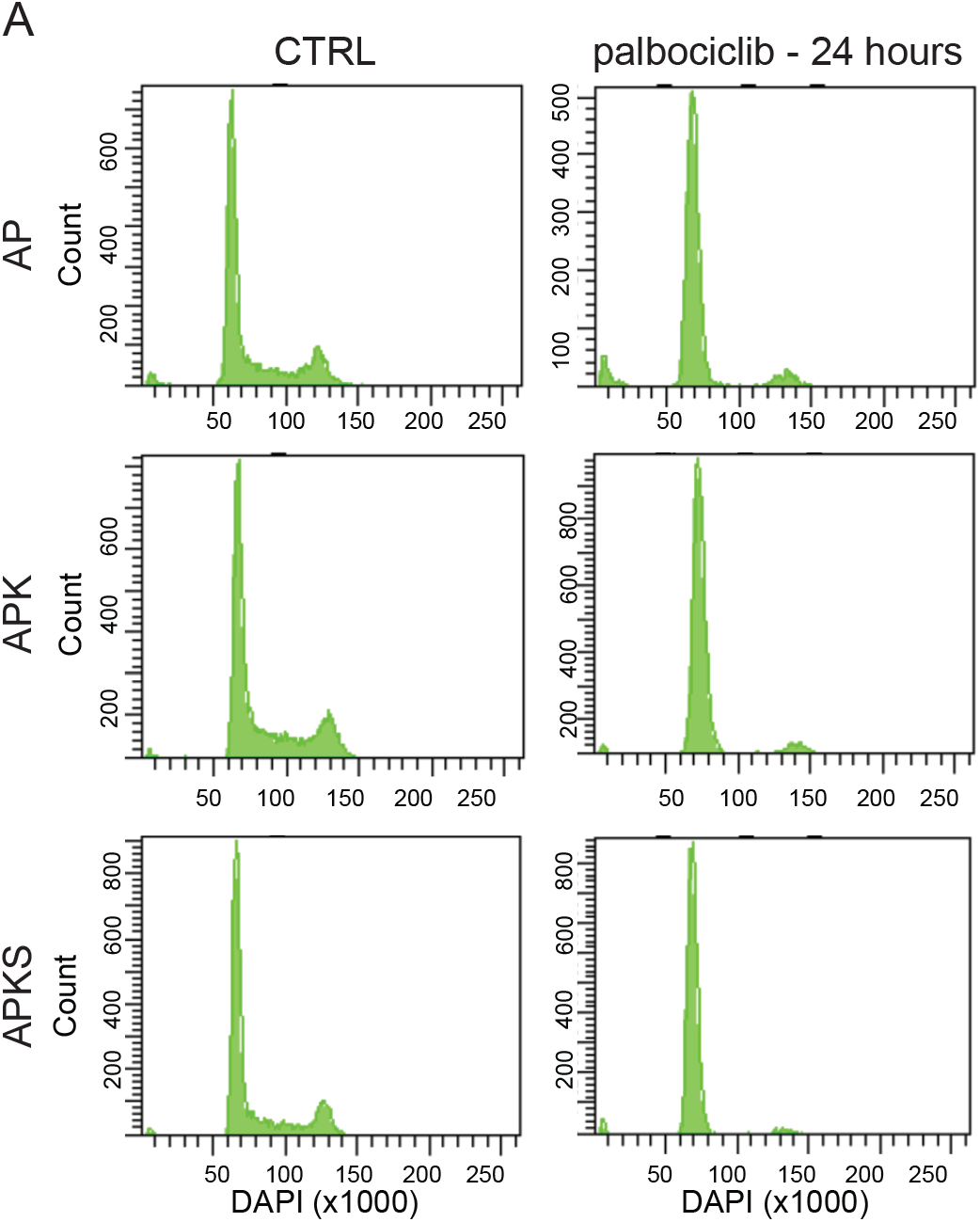
G1 arrest in AP, APK and APKS organoids by palbociclib treatment. **A)** Flow cytometry cell cycle profiles of AP, APK and APKS organoids stained with DAPI and treated with palbocilib for 24 hours.

**Figure S3.**
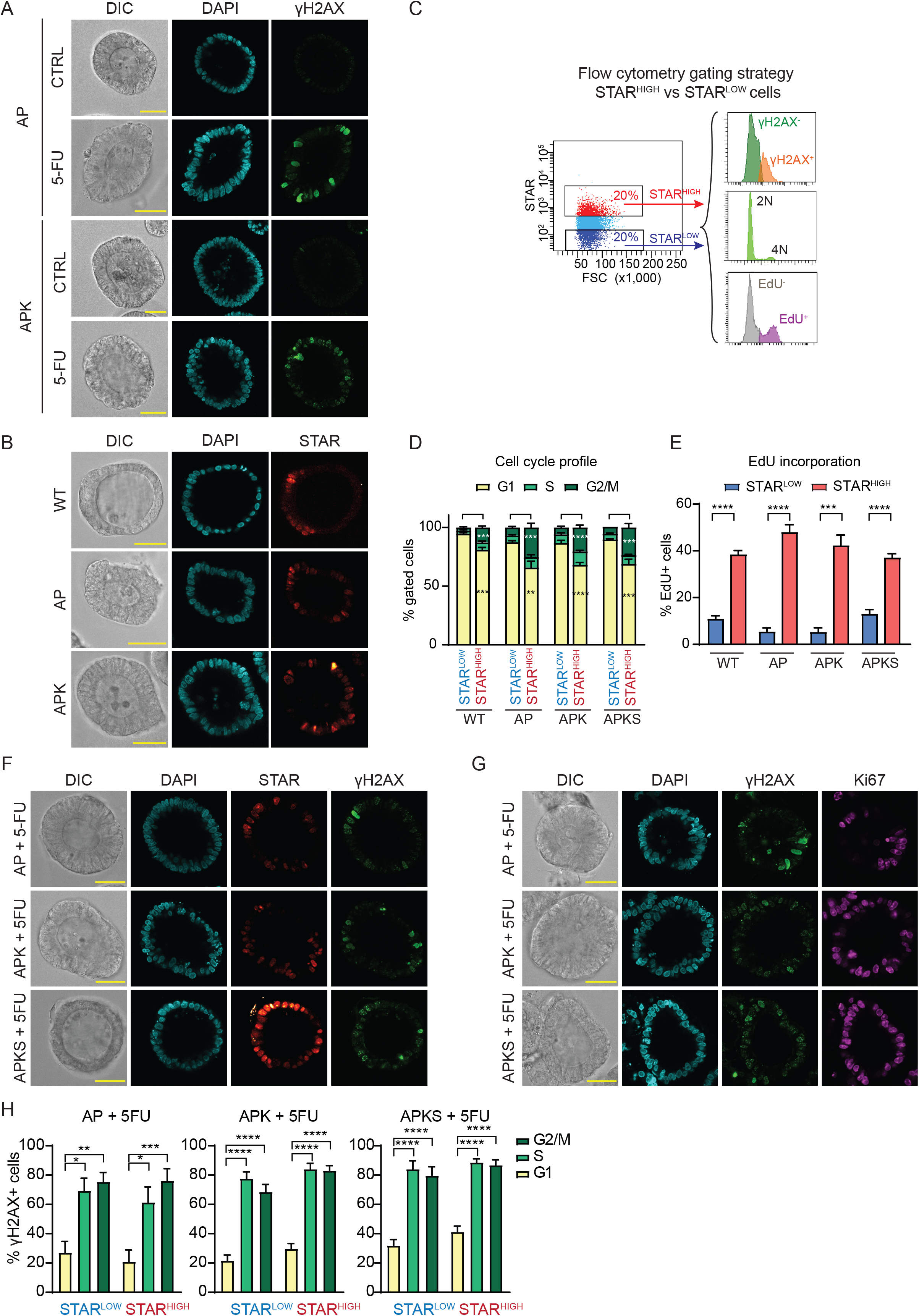
5-FU induces DNA damage in proliferating cells. **A)** Representative images of AP and APK organoids treated with 5-FU for 48 hours and stained with anti-γH2AX and DAPI (scale bar = 50 μm). **B)** Representative images of WT, AP and APK organoids transduced with the stem cell reporter STAR and stained with DAPI (scale bar = 50 μm). **C)** Flow cytometry gating strategy for DNA damage, cell cycle profile and EdU incorporation analysis in STAR^+^ vs STAR^-^ populations. **D)** Cell cycle profile determined by flow cytometry in STAR^+^ vs STAR^-^ cells of WT and CRC organoids (mean ± SEM, n = 4-6, unpaired t-tests). **E)** EdU incorporation analysis by flow cytometry in STAR^-^ vs STAR+ cells in WT and CRC organoids (mean ± SEM, n = 4-5, one-way ANOVA, Sidak’s multiple comparisons test). **F)** Representative images of AP, APK and APKS organoids transduced with the stem cell reporter STAR treated with 5-FU for 48 hours and stained with anti-γH2AX and DAPI (scale bar = 50 μm). **G)** Representative images of AP, APK and APKS organoids treated with 5-FU for 48 hours and stained with anti-KI67 and DAPI (scale bar = 50 μm). **H)** Detection of γH2AX^+^ cells by flow cytometry G1, S and G2/M cells in the STAR^low^ and STAR^high^ population of CRC organoids treated with 5-FU for 48 hours. Percentages γH2AX^+^ cells are normalized per specific cell type (mean ± SEM, AP: n = 5, APK: n = 7, APKS n = 6, one-way ANOVA, Sidak’s multiple comparisons test). ** p < 0.01, *** p < 0.001, **** p < 0.0001.

**Figure S4.**
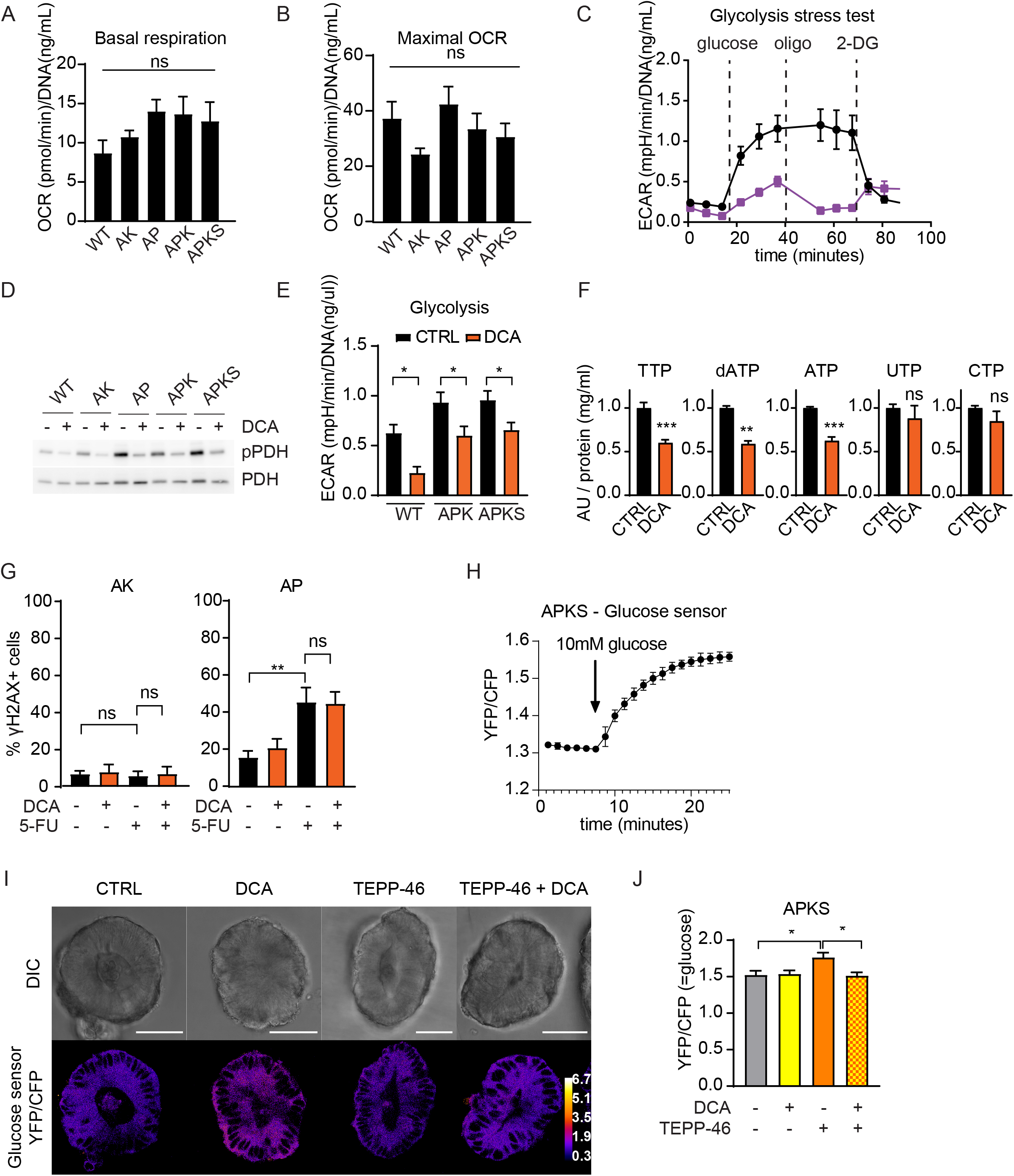
Respiratory phenotype of CRC organoids and metabolic analysis of 2-DG, DCA and TEPP-46 treatments. **A, B).** Basal oxygen consumption rate (OCR) and maximal respiration of WT and CRC organoids determined by mitochondrial stress test by Seahorse XF analysis (mean ± SEM, n = 5, one-way ANOVA). **C)** Determination of extracellular acidification rate (ECAR) during a Seahorse XF glycolysis stress test of APKS organoids treated with 2-DG for 24 hours (mean ± SEM, 5 technical replicates, representative for n = 3). **D)** Western blot detection of (p)PDH of WT and CRC organoids treated with DCA for 24 hours (blot representative for n = 3). **E)** Extracellular acidification rate (ECAR) of WT, APK and APKS organoids treated with DCA for 24 hours, determined by a glycolysis stress test by Seahorse XF analysis (mean ± SEM, n = 4-7, one-way ANOVA, Sidak’s multiple comparisons test). **F)** Detection of (deoxy)nucleotide triphosphates by metabolomics of APKS organoids treated with DCA for 20 hours (mean ± SEM, n=2 with 3 technical replicates each (dATP only detected in 1 experiment), one-sample t-test. **G)** Quantification of cells with DNA damage by flow cytometry of AK and AP organoids treated with 5-FU for 48 hours. DCA treatment started 20 hours before 5-FU treatment (mean ± SEM, WT: n = 3-6, AP: one-way ANOVA, Sidak’s multiple comparisons test, AK: Kruskal-Wallis test, Dunn’s multiple comparisons test). **H)** YFP/CFP ratio APKS organoids transduced with a FRET-glucose sensor treated with 10 mM glucose as validation of the glucose sensor (mean ± SEM, 5 technical replicates, YFP/CFP ratio resembles glucose concentration). **I, J)** Representative images of APKS organoids transduced with the FRET-glucose sensor treated with DCA and TEPP-46 for 24 hours (scale bar = 50 μm) (G), and quantification of these images (mean ± SEM, 17-22 organoids from 4 independent experiments, Kruskal-Wallis test, Dunn’s multiple comparisons test) (H) (YFP/CFP ratio resembles glucose concentration). ns: non significant, * p < 0.05

**Figure S5.**
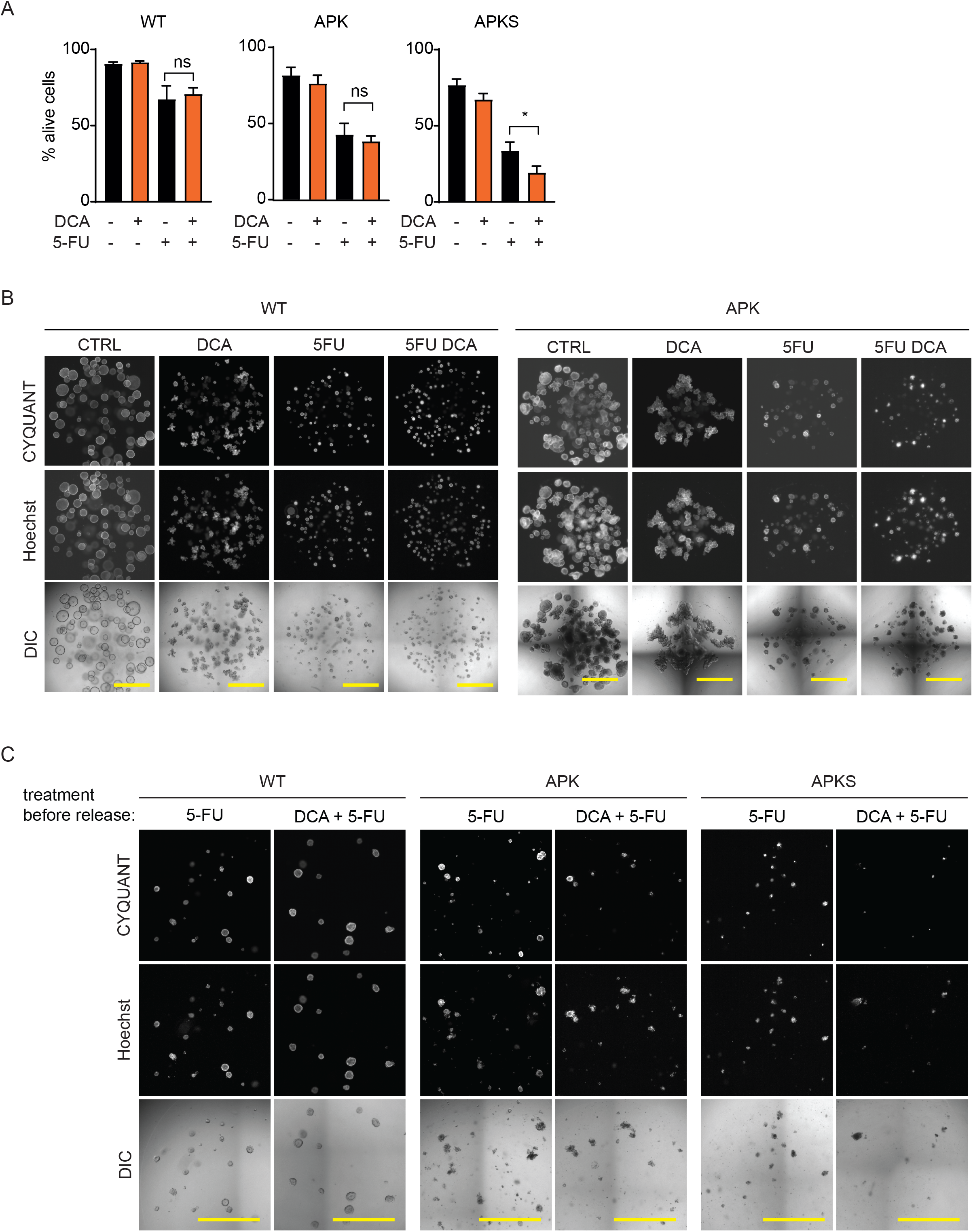
Targeting the Warburg effect by DCA improves 5-FU-induced cytotoxicity specifically in CRC organoids. **A)** Cell viability analysis of WT and CRC tumor organoids by flow cytometry to distinguish alive (DAPI^-^) from death cells (DAPI^+^) upon 7 days of 5-FU treatment. DCA treatment started 20 hours before 5-FU administration (mean ± SEM, n = 4-7, one-way ANOVA, Sidak’s multiple comparisons test). **B)** Representative images of WT and APK organoids, stained with CYQUANT (alive) and Hoechst (total), treated with 5-FU for 7 days. DCA treatment started 20 hours before 5-FU treatment (scale bar = 1 mm). **D)** Representative images of WT, APK and APKS organoids 4 days after replating, upon a 48 hour-5-FU (50 μM) treatment (with or without DCA treatment that started 20 hours before 5-FU administration), followed by 7 days of recovery time. Organoids were stained with CYQUANT (alive) and hoechst (total) (scale bar = 1 mm). ns: non significant, * p < 0.05

